# A meta-analysis and experiment assessing phage-based FMDV vaccine

**DOI:** 10.1101/2020.09.18.303107

**Authors:** Peng Wu, Ningning Yang, Yueli Wang, Mingguo Xu, Yunfeng Zhang, Chuangfu Chen

**Affiliations:** College of Animal Sciences, Shihezi University, Shihezi, Xinjiang, China; School of Animal Science and Technology, Shihezi University, Shihezi, Xinjiang, China; Co-Innovation Center for Zoonotic Infectious Diseases in the western region, Shihezi, Xinjiang, China; Key Laboratory of Control and Prevention of Animal Disease, Xinjiang Production & Construction Corps, Shihezi, Xinjiang, China; State Key Laboratory of Sheep Genetic Improvement and Healthy Production/Xinjiang Academy of Agricultural and Reclamation Sciences, Xinjiang, China

## Abstract

Foot-and-mouth disease (FMD) is a pathological disease caused by the foot-and-mouth disease virus (FMDV), which mainly affects cloven-hoofed animals. This study was conducted to a meta-analysis and experiment on the effect of bacteriophages used in the development of FMDV vaccines. A systematic search was conducted for the collection of the protection effect for the phage-based FMDV vaccine using sensitive search strategies. The extracted data were analyzed using Rev-Man 5.4 software. This experiment used the T7 phage to express the capsid protein VP1 of the OHM-02 strain, and the recombinant VP1 phage was termed OHM-T7. Antibodies and cytokines levels were assessed after immunizing BALB/C mice with OHM-T7. The results showed that a total of 115 articles were retrieved, and 4 of them met the inclusion criteria. There was no heterogeneity with I^2^ = 0%, 20% or 43%. We used a fixed-effect model for meta-analysis, and the results showed a protective effect on FMDV between the phage group and control group (*P*<0.01) and between FMDV group and control group (*P*<0.01). Furthermore, when the phage group was compared to the FMDV group, there was also no significant difference (*P*>0.05). After successfully obtained the ohm-t7 strain and immunized the mice, it could induce high levels of IFN-γ levels in mice with little effect on IL-4 levels. OHM-T7 could be used to detect antibodies produced by mice immunized with different FMDV antigens and produce high levels of anti-FMD antibodies. In summary, these results showed the potential of phage-based FMDV vaccines in FMDV prevention.

## Introduction

Foot-and-mouth disease (FMD) is a pathological disease caused by the foot-and-mouth disease virus (FMDV), which affects cloven-hoofed animals [1,2]. FMD causes serious economic and social problems, and is listed as a legally reportable disease by the World Organization for Animal Health (OIE) [3]. The virus has no capsule and has a diameter of about 30 nm. FMDV can infect cloven-hoofed animals, including pigs, sheep, goats, cattle and diverse wildlife species, and remains a major threat to the livestock industry worldwide [4]. Cattle could spread FMDV to pigs [5]. O-type FMDV is the most popular subtype around the globe. Currently, there is no vaccine that could protect animals from all serotypes [6]. Therefore, new vaccines are urgently needed to prevent the spread of FMD [7]. The capsid protein VP1 of FMDV is a sequence-dependent epitope, is the main antigenic site of FMDV, and can induce neutralizing antibodies.

Microphage (Phage) has been researched for decades [8]. Smith confirmed in 1985 that exogenous DNA could be inserted into the filamentous phage gene III and fused to the pIII protein [9]. The phage display technology inserts a DNA sequence into the phage coat protein’s structural gene, allowing the foreign gene to be expressed along with the coat protein [10-12]. Studies have shown that phages can be used to simulate viral epitopes [13,14]. The asymmetry of phage can enhance the immune response of helper T cell-1 and cause CD4^+^T cells to secrete cytokines [15,16]. The current phage display technology can insert DNA sequences into the structural genes of phage coat proteins so that foreign genes and coat proteins can be expressed together [10,12,17]. Phage display technology has been shown to increase the stability and immunogenicity of the antigen [18-20]. The phage display system is an ideal B cell epitope display vector and recombinant virus-like particles (VLPs) [21]. Among them, VLPs were found to be as immunogenic as the native virus, and the main reason for this is that the capsids are more heat-labile [22]. Therefore, phage vaccine, as a particulate antigen, can be quickly taken up by antigen presenting cells (APC). VLPs showed high immunogenicity and are easily recognized by the immune system [23]. Phage particles can also induce a strong cellular immune response [15]. Related studies have found that the peptides displayed by T7 phage bind closely to FMD serum [24]. To develop an effective FMD vaccine, we here conducted a meta-analysis and used a phage to display the FMD VP1 protein.

## Materials and Methods

### Literature search strategy

The literature retrieved in this meta-analysis was evaluated before March 2020 by two researchers. The National Library of Medicine (Medline via PubMed), Embase, China National Knowledge Infrastructure (CNKI), and Wan fang DATA were searched for phage-based FMDV vaccines, using the keywords “FMDV,” “phage,” and “vaccine.” Inclusion and exclusion criteria We used the following inclusion criteria: ① Published documents included Chinese and English literature. ② FMDV was expressed by the phage vector. ③ The protective effect of FMDV was evaluated in terms of the lethal dose. We used the following exclusion criteria: ① Method dissimilar to FMDV vaccine. ② The carrier was not a phage. ③ The results did not provide the necessary basic data.

### Data extraction tired

The two researchers conducted a preliminary screening by reading the titles and abstracts of the previous studies, then read the full text and made their selection according to the inclusion and exclusion criteria. If there were differences of opinion, we had already discussed and solved them. We independently extracted the data. The data extracted included the first author, publication time, events, and total number of animals in the trials.

### Analysis of extracted data

The database was developed using Excel. RevMan 5.4 software was used in this meta-analysis to perform the statistical analyses. The fixed-effect model was used for meta-analysis to calculate the odds ratios (OR), together with a 95% confidence interval (CI) for dichotomous results. ORs were used to evaluate the difference in immunogenicity between the two groups. A OR=1 indicates that data is of no worth. OR above 1.0 correspond to an effect favoring vaccination. Statistical heterogeneity between the studies was assessed using the I^2^ statistic and Q statistic. *P*≥0.05 or I^2^≤50% indicated that the trials were free of heterogeneity, and a fixed-effect model was used to perform the meta-analysis. I^2^>50% led us to consider a random-effect model to perform the meta-analysis. Where applicable, we presented results from individual trials and the common effect estimate in a forest plot. Squares indicate individual study odds ratios together with their 95% CI indicated as bars. Sensitivity analysis was performed using the difference of the combined values of the model effects, and the funnel plot method was used to evaluate the publication bias of the included works.

### Evaluation of the quality of evidence

All of the included studies were animal experiments, and animal experiments may be the highest level of evidence.

### Construction of the recombinant vector

The VP1 gene plasmid of OMH-02 was doubles digested using *EcoR* I and *Hind* III (TaKaRa, Dalian, China). In the control group, no enzyme was added; in the negative control group, the enzyme was substituted by an equal volume of water. All samples were incubated in a water bath at 37°C for 4 h. The digested product was subjected to 1% agarose gel electrophoresis. The target band was excised and purified with a DNA gel recovery kit (TaKaRa, Dalian, China), according to the manufacturer’s instructions. The extracted fragment of interest was ligated into the T7Select® vector in a reaction (Merck KGaA, Darmstadt, Germany). The sample was added to a 1.5 mL EP tube, gently pipetted up and down, incubated at 16°C for 16 h, and stored at 4°C.

### Phage packaging and plaque assay

The T7Select® package extract was thawed on ice. Then, 5 μL of the extract was added to 5 μL of the ligation reaction. The mixture was reacted at 22°C for 2 h, and 270 μL of TB medium was added. The phage was supplemented to *E*.*coli* BLT5403 in the logarithmic growth phase, and cultured at 37°C for 3 h. *E*.*coli* BLT5403 strain was inoculated in M9TB medium and incubated at 37°C with shaking to OD_600_ =1.0. Molten agarose was incubated in a water bath at 50°C. A 10^3^-10^6^ diluted sample was prepared with sterile TB medium as a diluent. Next, 250 microliters of BLT5403 was added to the EP tube, followed by addition of 100 μL of phage dilution and 3 mL of top agarose. The mixture was poured into an agar plate, inverted and incubated at 37°C for 4 h. The plaques were counted, and phage titers were determined.

### Identification of OHM-T7

A single plaque was scraped off using a pipette tip and heated at 65°C for 10 min. The sample was cooled to room temperature and centrifuged at 14,000 × g for 3 min. Primers were: T7 Select-F 5′-GGAGCTGTCGTATTCCAGTC-3′and T7 Select-R 5′-AACCCCTCAAGACCCGTTTA-3′, SDS-PAGE was run with 12% separation and 5% stacking gels. Staining was performed with Coomassie Brilliant Blue staining solution. OHM-T7 screening

A ninety-six-well ELISA plate was rinsed 3 times with deionized water. The Bovine FMD serum samples were diluted 30-fold with the ELISA coating solution, and 100 μL was added per well of the ELISA plate, which was incubated overnight at 4°C. After two washes with PBST, 200 μL/well of 5% skim milk powder was added and incubated at 37°C for 2 h. The plates were washed 3 times with PBST, and OHM-T7 was added for 2 h at 37°C. The cells were washed 3 times with PBST, and BLT5403 cells in the logarithmic growth phase were added to the wells and cultured in a 37°C incubator for 1 h. The bacterial suspension was aspirated and added to 20 mL of BLT5403 culture in the logarithmic growth phase, cultured at 37°C for 2 h. The library was preserved, and the product was used for subsequent screening. The above experiment was repeated 3 times.

### Immunization of mice

Twenty-four female BALB/C mice were purchased from Huaxing Laboratory Animal Farm (Huiji Distract, Zhengzhou, China). All the experimental procedures involving animals were approved by the Animal Experimental Ethical Committee Form of the First Affiliated Hospital of Medical College, Shihezi University (No. A 280-163-01). Female BALB/C mice were divided into the NaCl group, prokaryotic group and phage group, with 8 mice in each group. In the NaCl group, 250 microliters of normal saline were injected. The OHM-02 VP1 group was injected with prokaryote-expressed FMD VP1 protein at 200 µg/mice in a volume of 250 microliters. The OHM-T7 group was injected with 250 μL of OHM-T7 at a titer of 4×10^11^ Colony-Forming Units (CFU). Serum was collected at 0, 14, 28, 42, 56, 70, 84 and 98 d, respectively, and antibody levels were measured by ELISA. The OHM-T7 titer was 4×10^11^ CFU, and the sample was diluted 200-fold with the ELISA coating solution. IL-4 and IFN-γ were detected using an ELISA kit (Solarbio, Beijing, China).

### Statistical Analysis

The results of ELISA were analyzed with the SPSS 17.0 software (SPSS, Inc. Chicago, IL, USA), and all other statistics were performed using the GraphPad Prism 6 software package (Monrovia, CA, USA). A P-value of < 0.01 was considered greatly significant, and a P-value of < 0.05 was considered significant. All experiments were independently performed at least three times.

## Results

### Identified study reports

As shown in Fig. 1, document retrieval and filtering. A total of 115 articles were searched from databases. After deleting 12 repeated articles and reading the title and abstract, a total of 23 articles met the inclusion criteria. In the included literature, a total of 4 articles were included for meta-analysis.

**Fig. 1.**
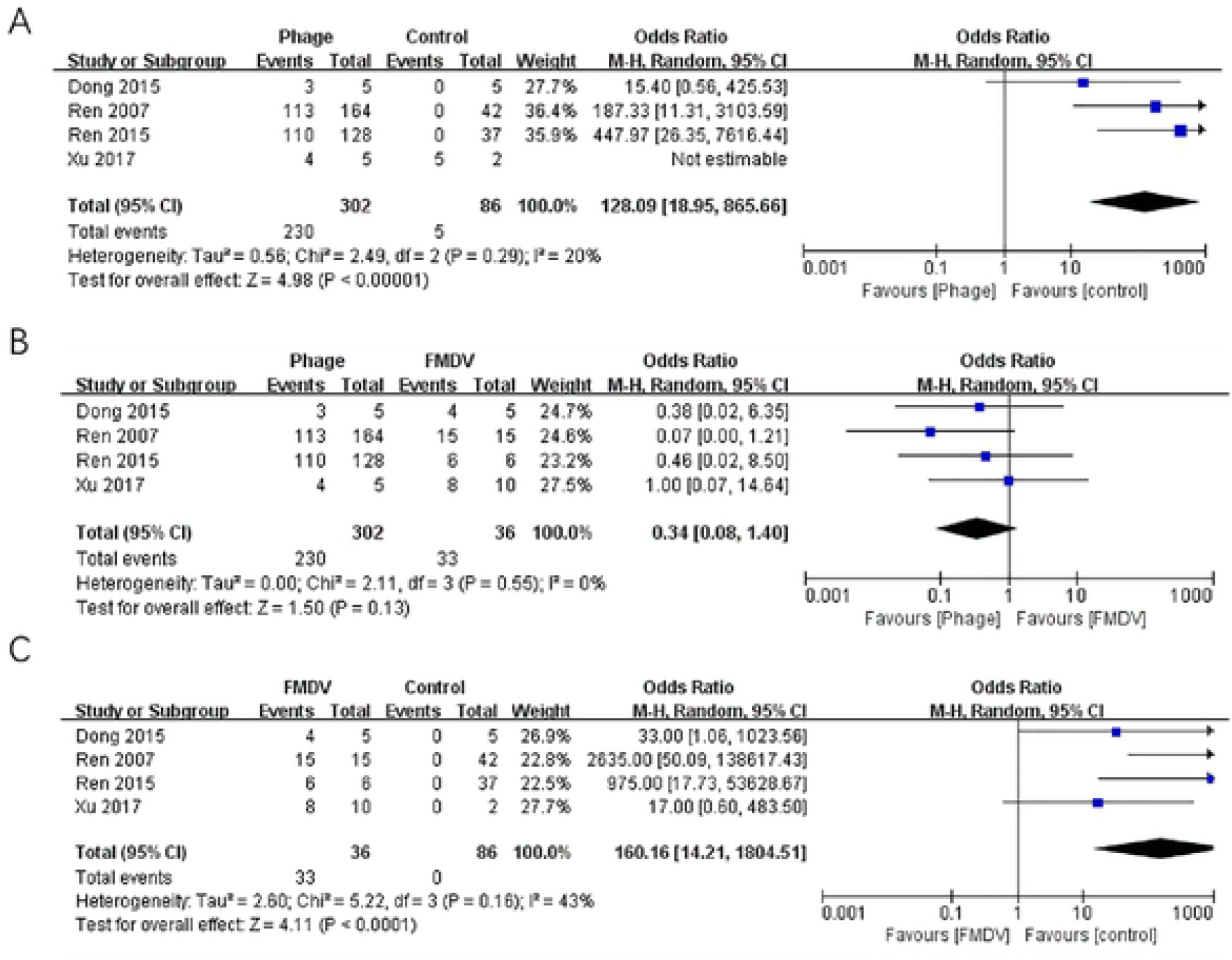
Flowchart of included and excluded trials.

### Characteristics of the reports

The characteristics of the included study are shown in Table 1. We also noted that all the studies were conducted between 2007 and 2017.

**Table 1.** Characteristics of eligible trials.

| Study |  | Phage Group |  | Control Group |  | FMDV Vaccine Group |  |
| --- | --- | --- | --- | --- | --- | --- | --- |
| Author | Year | Events | Total | Events | Total | Events | Total |
| Dong | 2015 | 3 | 5 | 0 | 5 | 4 | 5 |
| Ren | 2007 | 113 | 164 | 0 | 42 | 15 | 15 |
| Ren | 2015 | 110 | 128 | 0 | 37 | 6 | 6 |
| Xu | 2017 | 4 | 5 | 0 | 2 | 8 | 10 |

### Meta-analysis

In order to solve the problem of poor test results caused by the small number of documents, we here adopted a combination of statistical values and Q testing. The comparison of immune effects between the control group and experimental group was analyzed by fixed-effect model (Fig. 2).

**Fig. 2.**
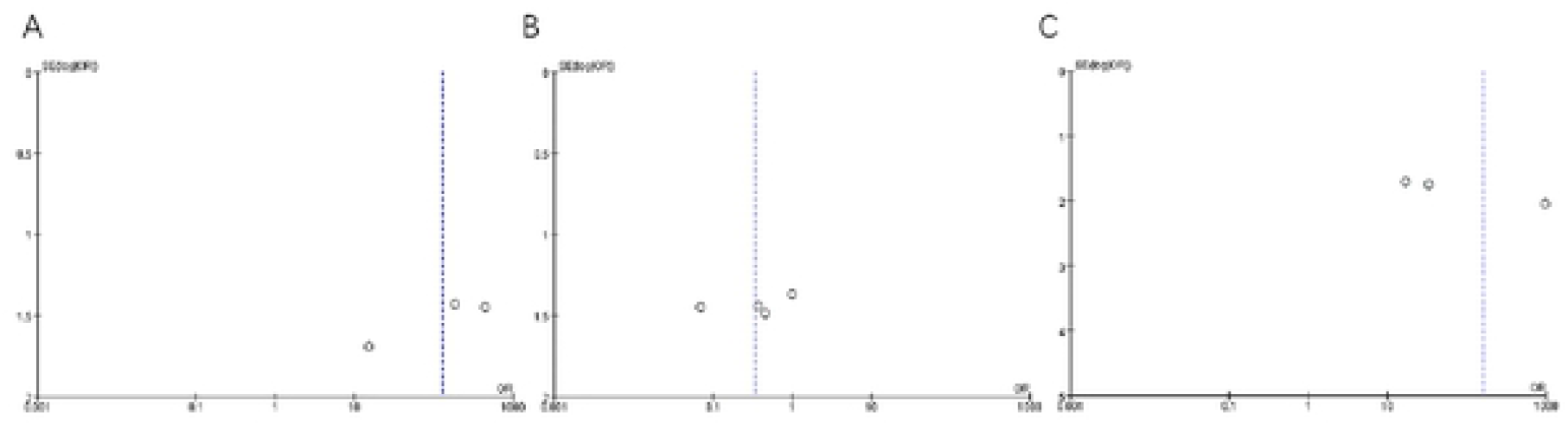
Forest plot of meta-analysis for ORR. There was no heterogeneity in this study with I^2^ = 20%, I^2^ = 0%, and I^2^ = 43%. We used a fixed-effect model for meta-analysis, and the results showed a protective effect on FMDV between the phage group and control group (MH = 128.09, 95% CI = 18.95, 865.66, *P*<0.01) (A) and between FMDV group and control group (MH = 160.16, 95% CI = 14.21, 1 804.51, *P*<0.01)(B). Furthermore, when the phage group was compared to the FMDV group, there was also no significant difference (MH = 0.34, 95% CI = 0.08, 1.40, *P*> 0.05) (C).

### Publication bias

The funnel chart method was used to control the publication bias of meta-analysis documents. By observing the funnel chart, we could see that although the pattern was not completely symmetrical, the data were still within the acceptable range (Fig. 3). The results showed that the published literature had less publication bias and met the requirements of this study.

**Fig. 3.**
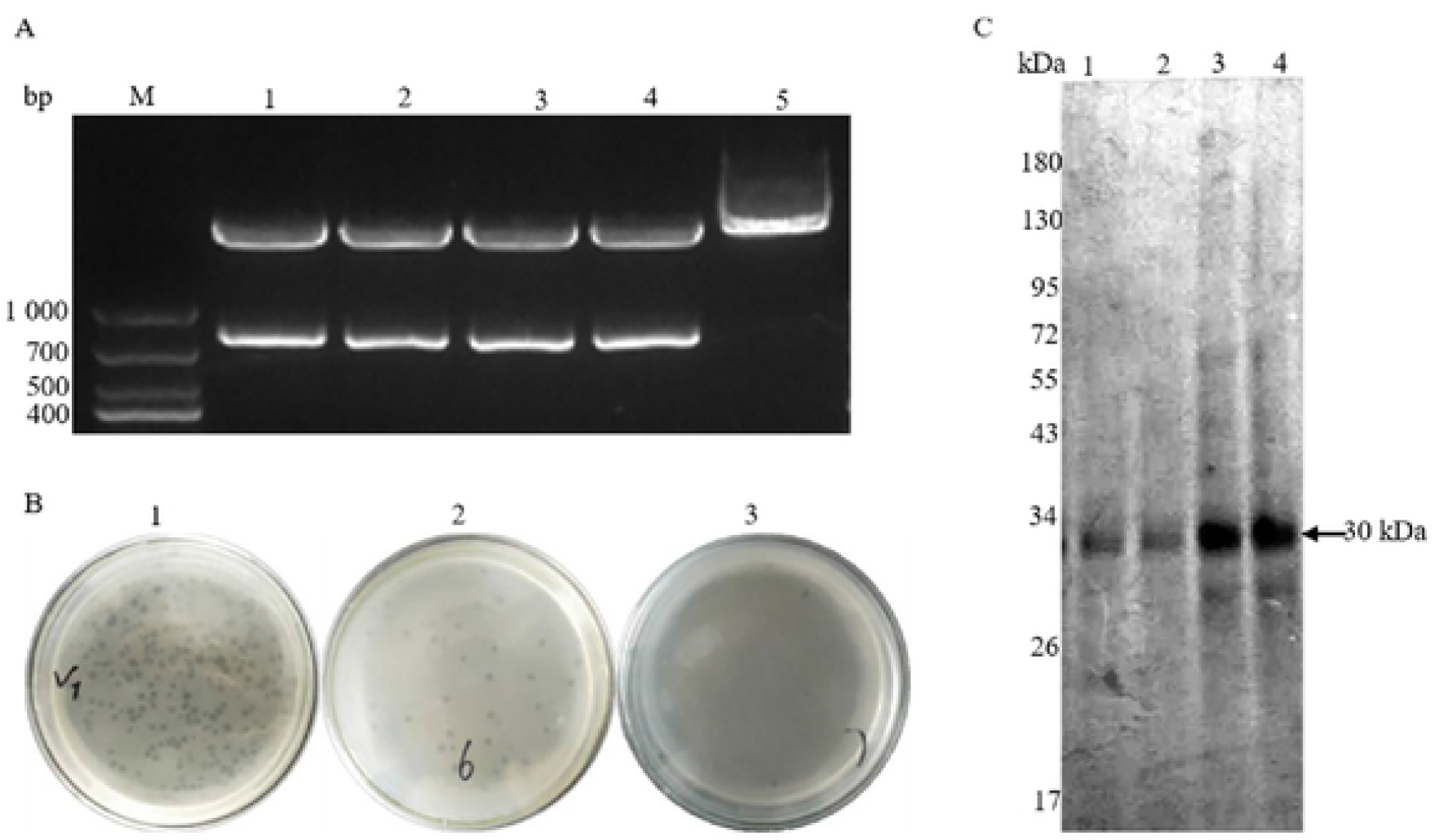
Funnel plot. (A) Phage group vs Control group; (B) Phage group vs FMDV group; (C) FMDV group vs Control group.

### Quality evaluation of evidence

All the experiments we selected evaluated the protective effect of the vaccine. All the experiments used animals for evaluation. The outcome of the protective trial was only survival or death, which was very different from the evaluation in humans.

### Construct and identification of the phage

After double digestion with *EcoR* I and *Hind* III, two clear bands were detected by 1% agarose gel electrophoresis. The target fragment was about 760 bp, which was consistent with the expected size (Fig. 4 A). The OHM-T7 titer was determined (Fig. 4 B). O5, O6 and O7 plates were 10^6^, 10^7^ and 10^8^ times diluted, respectively. Too many plaques grew on plate 5, and plates 6 and 7 had 40 and 4 plaques, respectively, indicating a titer of (4×108+4 ×108)/2=4×10^8^. Protein gel electrophoresis showed that OHM-T7 bands could be clearly observed at about 30 kDa, which proved that the phage was successfully expressed and purified (Fig. 4 C).

**Fig. 4.**
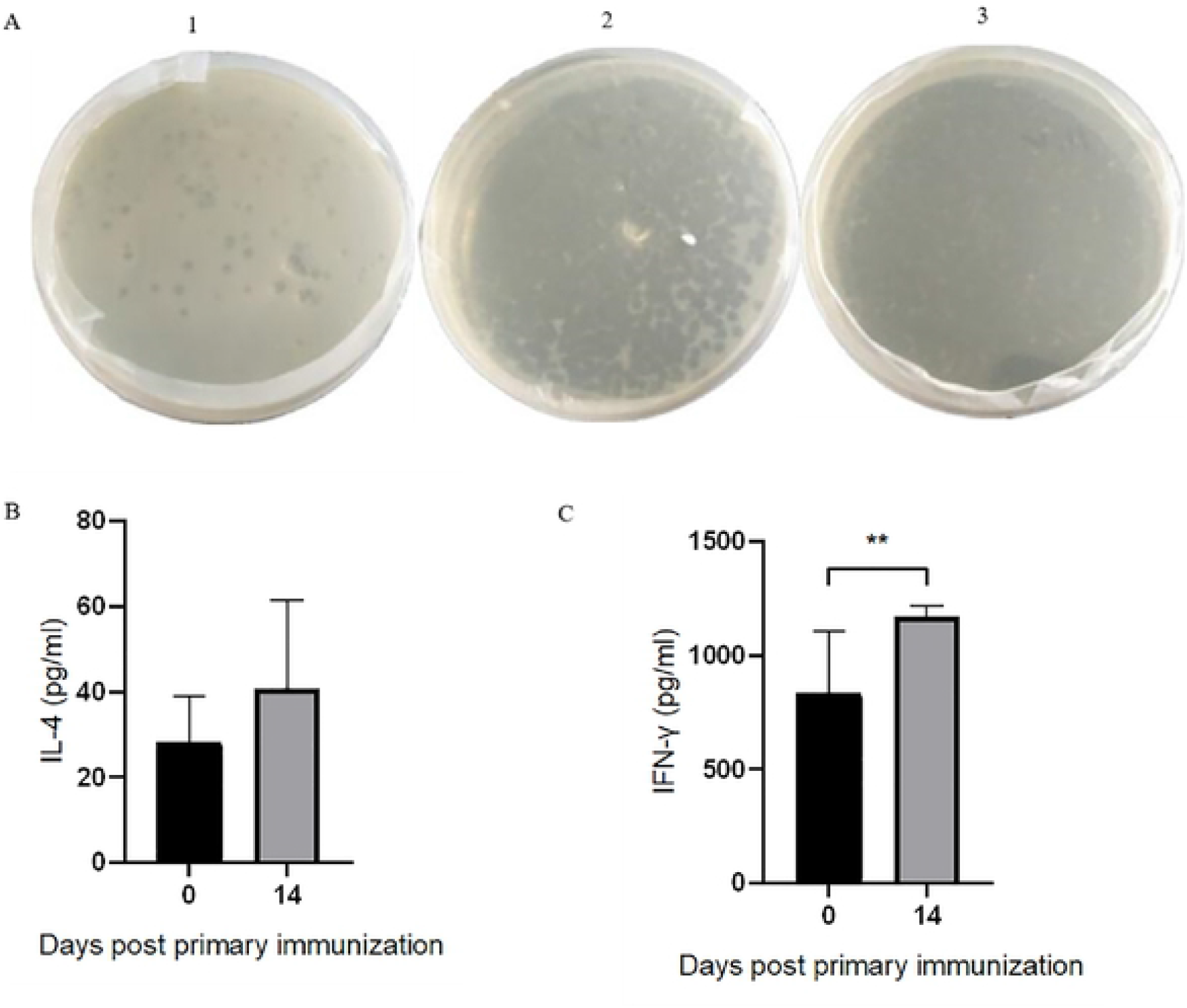
Construction of OHM-T7. (A) OHM-02 VP1 double restriction. M: DNA Marker 1000; Lanes 1-4: Double enzyme digestion positive clones; Lane 5: control group. (B) OHM-T7 plaque assessment; (C) OHM-T7 SDS-PAGE. Lanes 1-2: AKT-T7 control; Lanes 3-4: OHM-T7.

### Screening

The first round of screening of OHM-T7 on the Figure 5 A-1 plate yielded a titer of 4×10^10^. The titers of the second (Fig. 5 A-2) and third (Fig. 5 A-3) rounds for OHM-T7 plate were too high. The results showed that the OHM-T7 was enriched, and that with fast passage ability was selected (Fig. 5 A). The OHM-T7 strain induced high levels of IFN-γ levels (*P*<0.01) in mice with little effect on IL-4 levels (*P*>0.05).

**Fig. 5.**
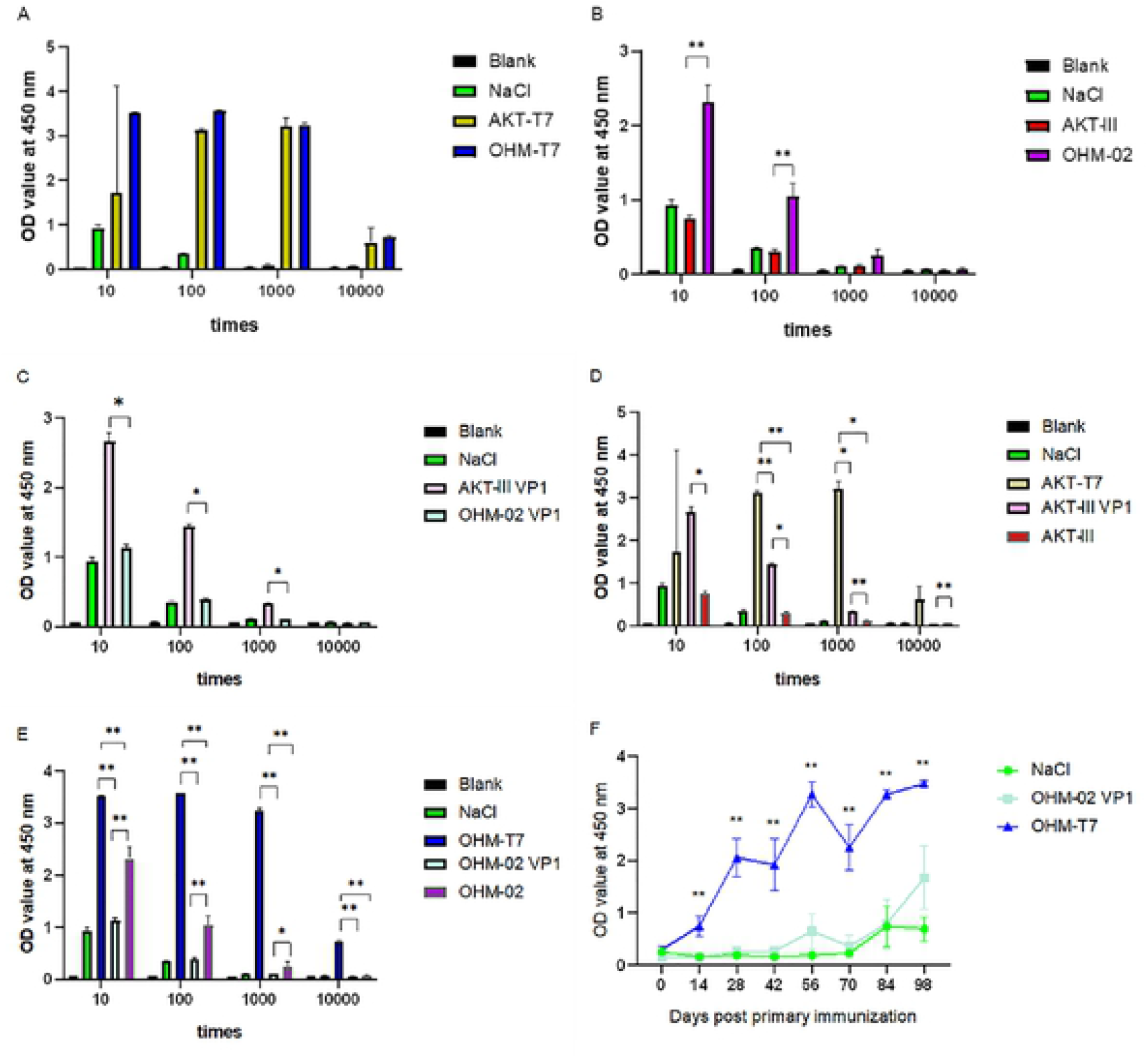
Immunogenicity of OHM-T7. (A) Reversal OHM-T7 reverse screening. (B) IL-4 results analyzed by ELISA kits. The OHM-T7 strain induced little effect on IL-4 levels in mice(*P*>0.05). (C) IFN-γ results analyzed by ELISA kits. The OHM-T7 strain induced high levels of IFN-γ levels in mice (*P*<0.01).

### Immunization of mice

The difference in anti-OHM-T7 antibody levels between 14 and 0 d was extremely significant (*P*<0.01). The results showed that OHM-T7 could quickly induce high levels of antibodies in mice. The difference in anti-OHM-T7 antibody levels between 98 and 0 d was also extremely significant (*P*<0.01). OHM-T7 antibody levels peaked at 98 d. The results showed that OHM-T7 could stimulate mice to produce anti-FMDV antibodies with high titers for a long time (Fig. 6 F). The level of IL-4 remained at a low level with no significant changes. The type II helper T cells (Th2) cells were less active after 14 d (Fig. 5B). Additionally, mice injected with the OHM-T7 strain had greatly improved levels of IFN-γ (Fig. 5C). An increase in IFN-γ indicates that type I helper T cells (Th1) are activated and IFN-γ is a hallmark cytokine of Th1 cells, which function is mainly to promote cellular immunity. This indicates that phage immunity was mainly induced by Th1 cells after 14 d. OHM-T7 could be used to detect antibodies produced by mice immunized with different FMDV antigens (Fig. 6).

**Figure 6.**
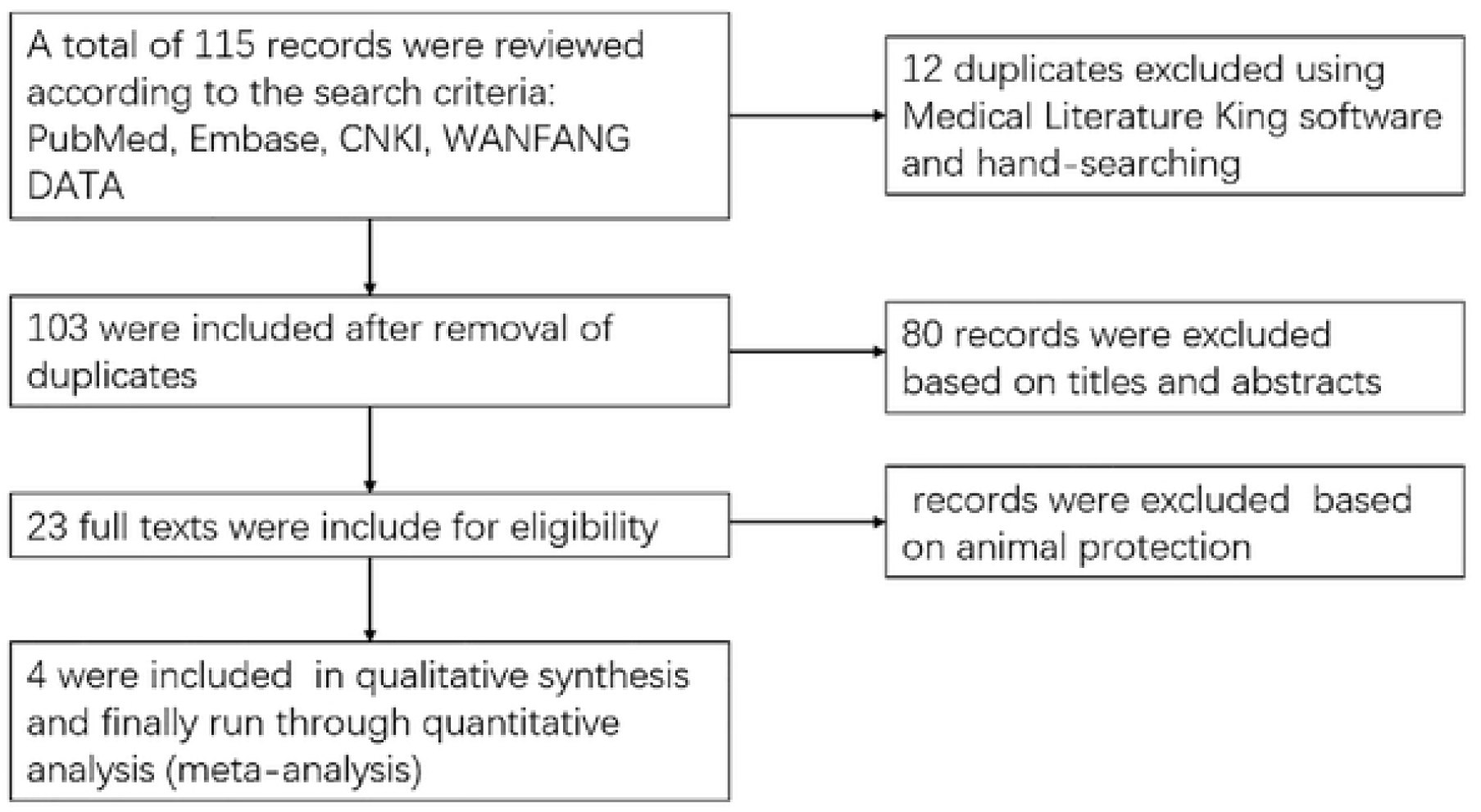
Detection of serum antibody levels in mice immunized with different FMDV antigens by OHM-T7. Blank, a blank control with nothing detected; NaCl, mouse serum immunized with 0.9% NaCl; AKT-T7, mouse serum immunized with the recombinant AKT-III VP1 phage; OHM-T7, mouse serum immunized with the recombinant OHM-02 VP1 phage; AKT-III VP1, mouse serum immunized with the prokaryotic expression the VP1 protein of the AKT-III virus; OHM-02, mouse serum immunized with the prokaryotic expression VP1 protein of the OHM-02 virus; AKT-III, mouse serum immunized with the AKT-III virus; OHM-02, mouse serum immunized with the OHM-02 virus. The data were analyzed by T-Text using excel software, * = *P* <0.05, ** = *P* <0.01.

## Discussion

This meta-analysis showed the promise of phage-based FMD vaccines. Bacteriophages have coexisted with humans and animals for a long time, and the safety of using them medically is widely recognized [25]. Recent studies have shown that phages play an important role in the mammalian immune system, and their interaction with mammalian immune cells is of great significance [26].

OHM-T7 were screened using immunopositively bovine serum samples, showing that the phage could bind to positive sera. After repeated screening, a highly specific OHM-T7 was obtained. The constructed phage library was reverse-screened, and OHM-T7 with high binding ability was selected. While obtaining a phage with a strong binding ability, phages with rapid propagation rates were also screened. Within 14 days, the phage could rapidly induce high levels of specific antibodies in the body. The current study only immunized the animals once, but after multiple blood collections the antibody levels remained high, demonstrating that the phage could stimulate the mouse body to maintain high antibody levels for a long time. Antibody levels were highest at 98 days, suggesting that antibody levels may increase. The antibody level of the AKT-T7 group was not different from that of the OHM-T7 group. It showed that the phage itself also caused the body to produce high levels of antibodies. The antibody level of the OHM-02 strain was found to be higher than that of the AKT-III group. It showed that OHM-T7 could specifically detect the O type antibody. It was very strange that the test result of AKT-III VP1 was higher than in the OHM-02 VP1 group. We attributed this to the instability of OHM-02 VP1 protein, and the results found in immunized animals were not ideal. The antibody level produced by the protein of the phage carrier was much higher than the prokaryotic expression. The phage vaccine produced much higher levels of antibodies than the inactivated virus vaccine.

Although we did not evaluate FMDV in animal protection experiments, the literature in the meta-analysis did include such experiments. In this way, our meta-analysis suggested that our recombinant phages may protect animals from FMDV. In 2009, the World Influenza Conference in Europe focused on vaccines for phage-derived virus-like particles as delivery vehicles [20]. Phage particle vaccines can also be administered via various immunization routes [26,27]. Under the conditions of large-scale breeding, good oral immunization effects are of great significance for the promotion and use of vaccines [28,29]. The use of a phage as a vector to display specific epitopes of different pathogens can induce a strong immune response in the body [30-33]. Therefore, the phage display technology has its unique value in the development of vaccines.

### Conclusion

The results of the present meta-analysis showed that the phage had protective effects on FMDV, and no difference was detected between the FMDV group and control with respect to this protective effect. The OHM-T7 was successfully constructed. OHM-T7 could be used to detect antibodies produced by mice immunized with different FMDV antigens and produce high levels of anti-FMD antibodies. This meta-analysis and experiment indicate the potential of phage-based FMDV vaccines in FMDV prevention.

## Acknowledgment

This research was funded by CX Collaborative Innovation 2011 Collaborative Innovation Special Project (Grant no. 0101-KC-0003).

## Author Contributions

Conceptualization: CC.

Data curation: PW, NY, MX, YZ.

Formal analysis: PW. NY.

Funding acquisition: CC.

Investigation: MX, YZ.

Methodology: PW, NY, YW.

Project administration: CC.

Resources: CC, PW.

Software: PW, NY.

Supervision: CC, PW.

Validation: CC.

Visualization: MX, YZ.

Writing – original draft: PW.

Writing – review & editing: PW, NY, Y.W.

## References

1. Niedbalski W, A. Kȩsy. Serological differentiation of animals infected and vaccinated against foot-and-mouth disease virus. Med. Weter. 2006; 62: 20–23. https://doi.org/10.1258/002367706775404417.

2. Ryan E, Mackay D, Donaldson A. Foot-and-mouth disease virus concentrations in products of animal origin. Transbound. Emerg. Dis. 2008; 55: 89–98. https://doi.org/10.1111/j.1865-1682.2007.01004.x.

3. Thomson GR, Vosloo W, Bastos ADS. Foot-and-mouth disease in Wildlife. Virus Res. 2003; 91: 145–61. https://doi.org/10.1016/S0168-1702(02)00263-0.

4. Lv J, Ding Y, Liu X, Pan L, Zhang Z, Zhou P, Zhang Y, Hu Y. Gene expression analysis of porcine whole blood cells infected with foot-and-mouth disease virus using high-throughput sequencing technology. Plos One 2018; 13: e0200081. https://doi.org/10.1371/journal.pone.0200081.

5. Mouton Laure, Dekker Aldo, Bleijenberg Meindert, et al. A foot-and-mouth disease SAT2 vaccine protects swine against experimental challenge with a homologous virus strain, irrespective of mild pathogenicity in this species. Vaccine; 2018, 36: 2020–2024. https://doi.org/10.1016/j.vaccine.2018.02.096.

6. Raza S, Siddique K, Rabbani M, et al. In silico analysis of four structural proteins of aphthovirus serotypes revealed significant B and T cell epitopes. Microb. Pathog. 2019; 254–262. https://doi.org/10.1016/j.micpath.2019.01.007.

7. Waters R, Ludi A B, Fowler V L, et al. Efficacy of a high-potency multivalent foot-and-mouth disease virus vaccine in cattle against heterologous challenge with a field virus from the emerging A/ASIA/G-VII lineage. Vaccine 2018; 1901–1907. https://doi.org/10.1016/j.vaccine.2018.02.016.

8. Comeau A M, Krisch H M. The Capsid of the T4 Phage Superfamily: The Evolution, Diversity, and Structure of Some of the Most Prevalent Proteins in the Biosphere. Molecular Biology & Evolution. 2008; 25: 1321–1332. https://doi.org/10.1093/molbev/msn080.

9. Smith G P. Filamentous Fusion Phage: Novel Expression Vectors that Display Cloned Antigens on the Virion Surface. Science 1985; 228: 1315–1317. https://doi.org/10.1126/science.4001944.

10. Bao Q, Li X, Han G, et al. Phage-based vaccines. Adv. Drug Deliv. Rev. 2019; 40–56. https://doi.org/10.1016/j.addr.2018.12.013.

11. Pande J, Szewczyk M M, Grover A K. Phage display: Concept, innovations, applications and future. Biotechnol. Adv. 2010; 849–858. https://doi.org/10.1016/j.biotechadv.2010.07.004.

12. Sidhu S S. Phage display in pharmaceutical biotechnology. Curr. Opin. Biotech. 2000; 11: 610–616. https://doi.org/10.1016/S0958-1669(00)00152-X.

13. De Andrade C Y, Yamanaka I B, Schlichta L S, et al. Physicochemical and immunological characterization of chitosan-coated bacteriophage nanoparticles for in vivo mycotoxin modeling. Carbohyd. Polym. 2018; 63–72. https://doi.org/10.1016/j.carbpol.2017.12.063.

14. Iniguez P, Zientara S. Selection of viral epitopes from phage display libraries. Application to diagnosis of equine arteritis virus. REV. MED. VET-TOULOUSE. 2001; 152: 363–371. https://doi.org/10.1038/srep21979.

15. Kolly R, Thiel M A, Herrmann T, et al. Monovalent antibody scFv fragments selected to modulate T-cell activation by inhibition of CD86–CD28 interaction. PROTEIN. ENG. DES. SEL. 2007; 20: 91–98. https://doi.org/10.1093/protein/gzl058.

16. Yang Q, Wang L, Lu D, et al. Prophylactic vaccination with phage-displayed epitope of C. albicans elicits protective immune responses against systemic candidiasis in C57BL/6 mice. Vaccine 2005; 23: 4088–4096. https://doi.org/10.1016/j.vaccine.2004.07.005.

17. Witherell G W, Gott J M, Uhlenbeck O C. Specific Interaction between RNA Phage Coat Proteins and RNA. Prog. Nucleic Acid Res. 1991; 40: 185–220. https://doi.org/10.1016/S0079-6603(08)60842-9.

18. Massis L M, Braga C J, Sbrogioalmeida M E, et al. Anti-flagellin antibody responses elicited in mice orally immunized with attenuated Salmonella enterica serovar Typhimurium vaccine strains. Mem. I. Oswaldo Cruz 2008; 103: 606–610. https://doi.org/10.1590/S0074-02762008000600017.

19. Larralde O G, Martinez R, Camacho F, et al. Identification of hepatitis A virus mimotopes by phage display, antigenicity and immunogenicity. J. Virol. Methods 2007; 140: 49–58. https://doi.org/10.1016/j.jviromet.2006.10.015.

20. Silman, Nigel J. World Influenza Congress Europe 2009. Expert Rev. Vaccines 2010; 9, 273–275. https://doi.org/10.1586/erv.10.10.

21. Li X, Meng X, Wang S, et al. Virus-like particles of recombinant PCV2b carrying FMDV-VP1 epitopes induce both anti-PCV and anti-FMDV antibody responses. Appl. Microbiol. Biotechnol. 2018; 102: 10541–10550. https://doi.org/10.1007/s00253-018-9361-2.

22. Ganji VK, Biswal JK, Lalzampuia H, Basagoudanavar SH, Saravanan P, Selvan RPT, et al. Mutation in the VP2 gene of P1-2A capsid protein increases the thermostability of virus-like particles of foot-and-mouth disease virus serotype O. Appl. Microbiol. Biotechnol. 2018; 102: 8883–8893. https://doi.org/10.1007/s00253-018-9278-9.

23. Samoylova T I, Norris M D, Samoylov A M, et al. Infective and inactivated filamentous phage as carriers for immunogenic peptides. J. Virol. Methods 2012; 183: 63–68. https://doi.org/10.1016/j.jviromet.2012.03.032.

24. Wong C L, Yong C Y, Muhamad A, et al. A 12-residue epitope displayed on phage T7 reacts strongly with antibodies against foot-and-mouth disease virus. Appl. Microbiol. Biotechnol. 2018; 102: 4131–4142. https://doi.org/10.1007/s00253-018-8921-9.

25. Aghebati-Maleki L, Bakhshinejad B, Baradaran B, et al. Phage display as a promising approach for vaccine development. J. Biomed. Sci. 2016; 23: 66–66. https://doi.org/10.1186/s12929-016-0285-9.

26. Górski Andrzej, Ryszard M, Jończyk-Matysiak Ewa, et al. Perspectives of Phage– Eukaryotic Cell Interactions to Control Epstein–Barr Virus Infections. Front. Microbiol. 2018; 630–630. https://doi.org/10.3389/fmicb.2018.00630.

27. Orourke J P, Peabody D S, Chackerian B. Affinity selection of epitope-based vaccines using a bacteriophage virus-like particle platform. Curr. Opin. Virol. 2015; 76–82. https://doi.org/10.1016/j.coviro.2015.03.005.

28. Zuercher A W, Miescher S M, Vogel M, et al. Oral anti-Ig E immunization with epitope-displaying phage. Eur. J. Immunol. 2015; 30: 128–135. https://doi.org/10.1002/1521-4141(200001)30:1<128::AID-IMMU128>3.0.CO;2-X.

29. Piekarowicz A, Klyz A, Majchrzak M, et al. Oral Immunization of Rabbits with S. enterica Typhimurium Expressing Neisseria gonorrhoeae Filamentous Phage Φ6 Induces Bactericidal Antibodies Against N. gonorrhoeae. Sci. Rep-Uk. 2016; 6: 22549–22549. https://doi.org/10.1038/srep22549.

30. Courtney B C, Williams K C, Schlager J J. A phage display vector with improved stability, applicability and ease of manipulation. Gene 1995; 165: 139–40. https://doi.org/10.1016/0378-1119(95)00526-C.

31. Jonas Kügler, Nieswandt S, Gerlach G F, et al. Identification of immunogenic polypeptides from a Mycoplasma hyopneumoniae genome library by phage display. Appl. Microbiol. Biotechnol. 2008; 80: 447–458. https://doi.org/10.1007/s00253-008-1576-1.

32. Krut O, Bekeredjian-Ding I. Contribution of the Immune Response to Phage Therapy. J. Immunol. 2018; 200: 3037–3044. https://doi.org/10.4049/jimmunol.1701745.

33. Hodyra-Stefaniak K, Miernikiewicz P, et al. Mammalian Host-Versus-Phage immune response determines phage fate in vivo. Sci. Rep-Uk. 2015; 5: 14802–14802. https://doi.org/10.1038/srep14802.

